# Super-resolution Fluorescence Microscopy Reveals Densely Packed EboV Membrane Glycoprotein Clusters in the Cell Plasma-membrane During Ebola Infection

**DOI:** 10.1101/2025.09.10.674168

**Authors:** Partha P. Mondal, Jiby M. Varghese, S Aravinth, Neeraj Pant, Souvik Saha

**Affiliations:** Department of Instrumentation and Applied Physics, Indian Institute of Science, Bangalore 560012, INDIA; Centre for Cryogenic Technology, Indian Institute of Science, Bangalore 560012, INDIA

## Abstract

The glycoprotein (GP) of Ebola virus plays a central role in the viral entry into host cells. The detailed study of Ebola glycoprotein in the cellular system is unknown, mainly due to the resolution limit (∼ *λ/*2) of optical fluorescence microscopes. To overcome this, we propose to use single-molecule localization microscopy (SMLM), which enables the study of the role of glycoproteins at both the single-molecule and ensemble levels. Accordingly, a photoactivable recombinant probe *Dendra*2 − *EboV Glyc* is developed by linking Ebola glycoprotein with Dendra2. The transient expression of photoactivable recombinant protein in NIH3T3 cells indicated the dominant presence of glycoprotein in the plasma membrane with regions of densely packed glycoproteins. Super-resolution single-molecule imaging gives direct proof of glycoprotein clusters in the plasma membrane. The corresponding single-molecule analysis using DBSCAN clustering algorithm reveals a dense packing of glycoprotein with an estimated average density of 5345 *mol/µm*^2^ compared to the density in other regions of the membrane. The number of glycoproteins is estimated to be as high as 642 molecules per cluster and clusters spread area of ≈ 0.1219 *µm*^2^ after 48 *hrs* post-transfection. Noting that glycoprotein is involved in cellular attachment, endosomal entry, and membrane fusion, the formation of glycoprotein clusters in the membrane is significant in the progression of Ebola pathogenesis. A drug capable of dispersing the glycoproteins clusters could be of therapeutic potential to lessen the intensity of infection and to slow down the disease progression.

**Statement of Significance:** The fact that powerful fluorescence microscopes (such as SMLM) are available is a boon to disease biology, specifically to study the function of viral proteins at the cellular level. Ebola, one of the nocturnal viral diseases, is a threat and calls for study at the single-molecule level to understand the dynamics (kinetics and ensemble) of glycoprotein during viral infection. In this respect, the super-resolution microscopy techniques may help reveal the biological mechanism behind the infection process at the single-molecule level. A photoactivable fluorescent probe (*Dendra*2 − *EboV Glyc*) containing the gene-of-interest (EboVGP) is developed, and transfection studies are carried out, showing densely packed glycoprotein clusters in the cell plasma membrane with specific density, number, and size. We believe that dispersing these glycoprotein protein clusters could be of therapeutic potential.

## I. INTRODUCTION

Ebola Virus (EboV) is a filovirus that belongs to a family of single-stranded negative-sense RNA viruses. Ebola virus disease is characterized by severe hemorrhagic fever with a fatality rate of 25 − 90% [41]. The genome of the Ebola virus is about 19 *kb* long, which encodes seven genes; nucleoprotein(NP), P protein (VP35), matrix protein (VP40), glycoprotein (GP), second nucleoprotein (VP30), protein associated with the envelope (VP24) and RNA-dependent RNA polymerase (L) [3] [4]. Specifically, the GP plays a central role in immune evasion and viral entry into cells [5] [7].

The glycoprotein is produced as precursor *GP*_0_, which is cleaved by furin into disulfide-bonded *GP*_1_ (a receptor-binding unit of size ∼ 130 kDa) and *GP*_2_ (a fusion subunit of ∼ 30 − 40 kDa that anchors the protein to the viral membrane). Both proteins are expressed on the surface of the virion [8]. Specifically, the *GP*_1_ heterodimer contains a receptor-binding domain and a heavily glycosylated mucin-like domain. The heavily glycosylated “glycan cap” and mucin-like domain shield the receptor-binding region from immune recognition. The fusion subunit, *GP*_2_, possesses a fusion peptide and heptad repeats that are necessary for virus-cell membrane fusion [10] [11]. Together, they form a trimeric spike on the viral envelope, similar to Human Immunodeficiency Virus (HIV) envelope glycoprotein or Influenza virus hemagglutinin (HA)[40]. *GP*_1_ binds to attachment factors on host cells such as C-type lectins (DC-SIGN, L-SIGN), TIM-1, and phosphatidyl serine receptors. This promotes initial virus docking, especially on dendritic cells and macrophages [43] [44]. 293T cell-based studies indicate that the entry of Ebola virus is primarily GP-mediated via macropinocytosis (a form of endocytosis involving engulfment of the virus into endosomes), although there are reports of endocytic pathways as well [12] [13] [14]. In late endosomes, cathepsins B and L cleave *GP*_1_, removing the mucin-like domain and glycan cap, exposing the receptor-binding site of *GP*_1_. Cleaved *GP*_1_ binds to Niemann–Pick C1 (NPC1), a cholesterol transporter in endosomal membranes[46]. NPC1 is considered the essential entry receptor for Ebola virus [45]. The Binding to NPC1 triggers conformational changes in *GP*_2_, driving fusion of viral and endosomal membranes. This releases the viral RNA genome into the cytoplasm, initiating infection.

The Ebola virus glycoprotein (EboVGP) is a key target for vaccines, neutralizing antibodies, and inhibitors that block viral entry, as it is the sole viral protein displayed on the surface of the virion [6]. EboVGP over expression studies in 293T cells resulted in cytotoxicity and down-regulation of several proteins involved in cellular adhesion [16]. The step-by-step structural dynamics of EboVGP during fusion remain poorly resolved. An in-depth study of the glycoprotein on cellular components may give a novel insight into Ebola virus pathogenesis [9]. This calls for a drastic change of approach and an urgent need to evaluate viral protein (EboVGP) at the single-molecule level.

Existing microscopy techniques (such as widefield, two-photon, and confocal) are limited by diffraction of light and at best can produce images with resolution, *r >* 1.2*λ/*2*NA* [1][2]. However, recent developments in single-molecule-based super-resolution microscopy with resolution *<< r* may play a major role in deciphering biological processes in a cellular system [20] [21] [22] [23] [24]. These techniques have matured in recent years with additional capability of specificity, single-molecule precision, and volume imaging [25] [28]. As a result, these techniques are finding application across the spectrum of biological sciences (from fundamental cell biology to virology) [25] [20] [26] [27] [28] [19]. Recent super-resolution studies include, dengue viral infection in cellular system [29] [30], HA clustering in influenza (type-A) infection [26] [31] and confocal microscopy study of the effective infection of dengue virus to host cell [32] [33] [34]. However, super-resolution studies elucidating the role of EboVGP, its spatial distribution, and kinetics in the plasma membrane are yet to be reported. This information may serve as the starting point for developing antiviral therapies, their efficacy, and restriction during viral infection.

In view of the fact that current evaluation and understanding are based on the analytical techniques and confocal studies [35] [36] [37], it becomes imperative to investigate the EboV viral infection at the very single molecule level. To do so, we developed a photoactivable probe (Dendra2-EboVGlyc) using standard molecular cloning methods, and carried out single-molecule-based super-resolution imaging studies to understand GP-mediated entry and its kinetics in the cell plasma membrane. In this report, we present the EboVGP mechanism (post 24 hrs of transfection) with single-molecule precision in a cellular system via transfection studies. In addition, we carried out time-lapse imaging of Dendra2-EboVGP protein in the cell plasma membrane. Compared to the studies based on diffraction-limited light microscopy, SMLM has the dual advantage of directly visualizing single molecules and their ensembles/clusters with super-resolution capabilities and carrying out single-molecule analysis. Moreover, time-lapse imaging enables visualization of the entire clustering process at varying time-points, giving details of its collective dynamics (single protein location, kinetics, and cluster formation), as well as its localization to specific organelles. The resultant cluster parameters (number of clusters, cluster area, density, and copy number) are determined, which give vital information related to cell physiology during the viral infection process. Based on the present investigation, the mechanism of EboVGP is visualized in the cell plasma membrane.

## II. RESULTS

Here, we explore the role of EboVGP at the plasma membrane during post-entry stages of Ebola virus infection. To carry out cellular studies, we have chosen to work with the mouse embryonic fibroblast cell line, NIH3T3. We constructed a recombinant photoactivable plasmid-DNA probe (Dendra2-EboVGlyc), which is used for cell transfection studies.

### A. In-vitro expression of photoactivable recombinant probe Dendra2-EboVGlyc

The recombinant probe (Dendra2-EboVGlyc) necessary for carrying out super-resolution studies is constructed by conjugating the protein-of-interest (EboVGP) with photoactivable protein (Dendra2). The presence of EboVGP in the Dendra2 backbone was confirmed by PCR and restriction digestion analysis (Fig. 1B). The expected PCR amplicon size, restriction enzyme digestion profile, and Sanger sequencing together confirmed the successful integration of the ORF encoding the recombinant Dendra2-EboV glycoprotein into the plasmid vector (Fig. 1A, B). PCR of recombinant probe Dendra2-EboVGlyc using EboVGP specific primers gave an amplicon around 2000 bp and restriction analysis using the restriction enzyme pair Sac1 and Bam H1 released a fragment of ∼ 2000 *bp* (see, Fig. 1B). The intactness of open reading frame is confirmed by the sequence analysis of Dendra2-EboVGlyc. The complete plasmid map of Dendra2-EboVGlyc is shown in Fig. 1A. The expression of protein is confirmed by confocal fluorescence microscopy (ex:em = 480 nm : 510 nm). Fig. 1C shows the overlayed images of the brightfield and fluorescence images for NIH3T3 cells. This isolates transfected cells (green) along with other cells that are not transfected, indicating the expression of photoactivable fluorescent protein glycoprotein (Dendra2-EboVGlyc). Note that the green fluorescence of the transfected cell results from the fluorescence emitted by non-activated proteins. Subsequently, the cells were fixed after 24 hrs of transfection for super-resolution imaging and single-molecule studies.

**FIG. 1:**
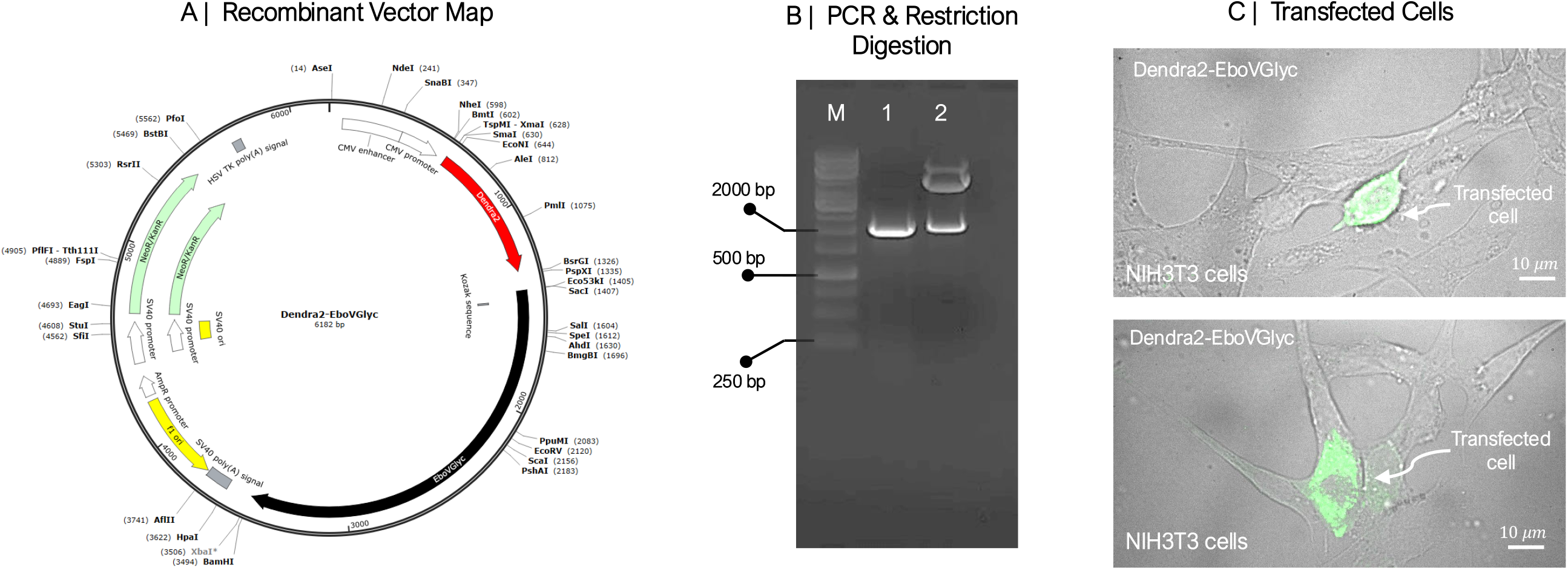
Confirmation and validation of recombinant probe Dendra2-EboVGlyc: (A) Vector map of recombinant plasmid *Dendra*2 − *EboV Glyc* showing restriction sites. Highlighted region with red and black indicates photoactivatable partner *Dendra*2 and protein of interest *EboV Glyc* respectively (B) Confirmation of recombinant probe by PCR and restriction digestion analysis showing amplification and fragment release around 2000 bp. (C) Overlay of fluorescent and transmission channel for the corresponding image shows clear distinction of transfected cell (green) from neighbouring non-transfected cells. Scale bar = 10 *µm*.

### B. Colocalization of EboVGlyc protein and Plasma-Membrane

To identify the activity of the recombinant protein, we have carried out colocalization studies. Accordingly, healthy NIH3T3 cells were first transfected with Dendra2-EboVGlyc plasmid DNA, and incubated for 24 hrs for the expression of the target protein (Dendra2-EboVGP). The colocalization of Dendra2-EboVGP at the plasma membrane was assessed by staining Dendra2-EboVGP plasmid transfected NIH3T3 cells with Lissamine Rhodamine B–Phosphatidylethanolamine (Liss-Rhod PE), a plasma membrane–specific dye. Fig. 2 shows the confocal images of a few chosen cells (Cells 1-3) along with the overlayed images. The red indicates cell membrane whereas, the green indicates the expressed Dendra2-EboVGP protein. It is noted that only the transfected cells show the colocalization (yellow color). This can be better seen in enlarged images of the transfected cells.

**FIG. 2:**
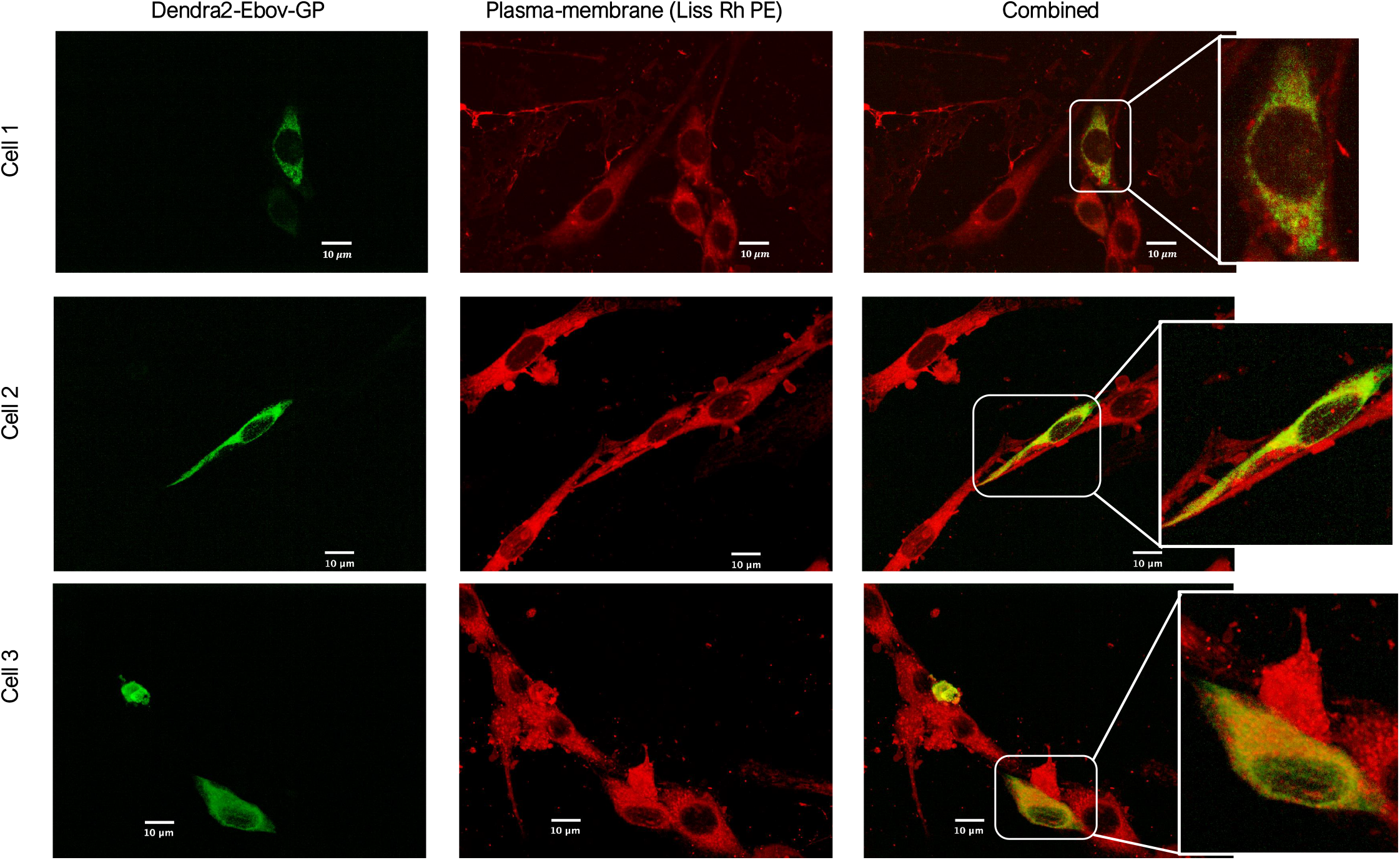
Localization of Dendra2-EboVGP in cell plasma membrane: Confocal images of photoactivatable Dendra2-EboVGP for few chosen cells (cell 1-3) obtained from multiple experiments, showing its localization to cell plasma-membrane. Here, membrane is labeled with Liss-Rhodamine B-Phosphatidylethanolamine (Liss-Rhod PE). Alongside, enlarged images are also shown.

In addition, colocalization analyses were performed at three post-transfection time points (24, 36, and 48 h) to evaluate the temporal dynamics of protein expression. NIH3T3 cells transfected with the Dendra2-EboVGP plasmid, were fixed and stained with the Liss-Rhod PE dye, at designated time interval (24, 36, and 48 h post-transfection). This approach enabled visualization of protein localization and dynamics relative to the plasma membrane overtime, post-transfection expression. Subsequently, all the samples are imaged using a confocal microscope, as shown in Fig. 3. Alongside, the transmission images are also displayed (panel 1). The expression of Dendra2-EboVGP is clearly evident in transfected cells (panel 2, green color), and the labeling of plasma-membrane by Liss-Rhod PE) is shown in panel 3 (red color). Visually, colocalization of transfected cells is evident from the overlayed images (combined, panel 4, yellow color). Subsequently, the images are subjected to colocalization analysis. Two different statistical tests (Overlay and Pearson) are carried out, and the coefficients are determined. A high value of the coefficients for the statistical colocalization parameters (*>* 0.8 for overlay; *>* 0.8 for Pearson) suggests the presence of the protein-of-interest (Dendra2-EboVGP) in the plasma membrane of the transfected cell.

**FIG. 3:**
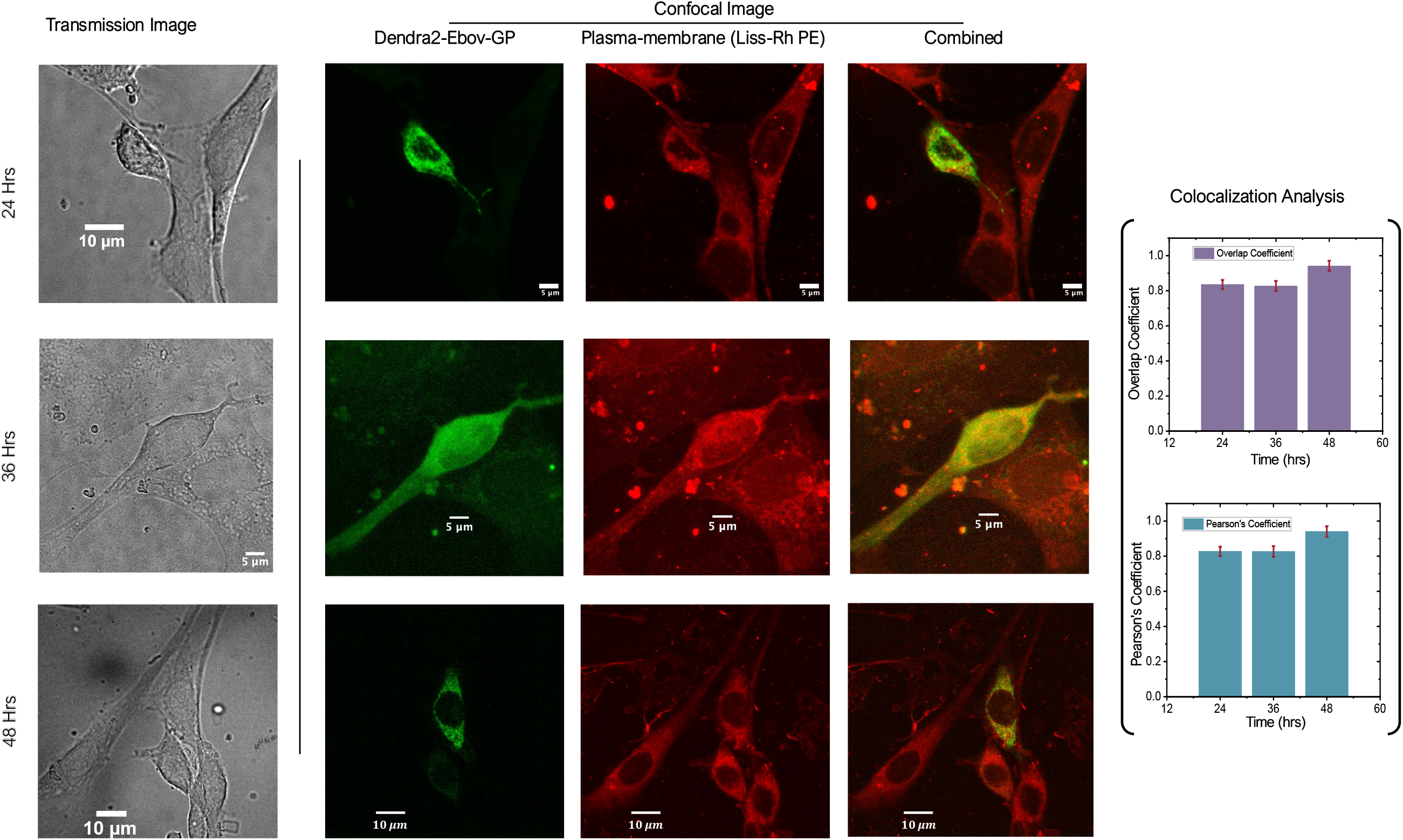
Colocalization analysis at varying time-points: Images of cells transfected with Dendra2-EboVGP and incubation for 24 hrs, 36 hrs and 48 hrs followed by staining with plasma membrane specific dye Liss-Rhod PE. Alongside transmission images are also shown. Colocalization analysis using statistical tests (Overlay and Pearson coeficient) were given in bar graph

### C. Super-resolution microscopy reveal Dendra-EboVGP clusters

To gain a better understanding of the single viral protein characteristics (location, kinetics, and its ensemble) in the plasma membrane, single-molecule-based super-resolution microscopy (SMLM) studies were carried out. Accordingly, healthy NIH3T3 cells were transfected with Dendra2-EboVGlyc plasmid DNA, and incubated for varying time points (24 hrs, 36 hrs, and 48 hrs). This allows direct visualization of the protein (Dendra2-EboVGP) expression and its location in the cell membrane. Subsequently, the cells were fixed and prepared for super-resolution imaging and single-molecule analysis. Fig. 4 shows the super-resolved images of transfected cells expressing the viral protein (Dendra2-EboVGP), along with the localization precision histogram. In addition, widefield fluorescence images (panel 1) and enlarged images of a few selected regions (dotted blue and orange boxes) are also displayed. It is evident that the viral protein Dendra2-EboVGP forms clusters. Moreover, the cluster increases in size over time and stabilizes, as evidenced by the cluster sizes at 48 hrs as compared to 24 hrs. This is impressive, as the observation of stable protein clusters in the cell plasma membrane over time may illustrate the biological mechanism of cluster formation during viral infection at a cellular level.

**FIG. 4:**
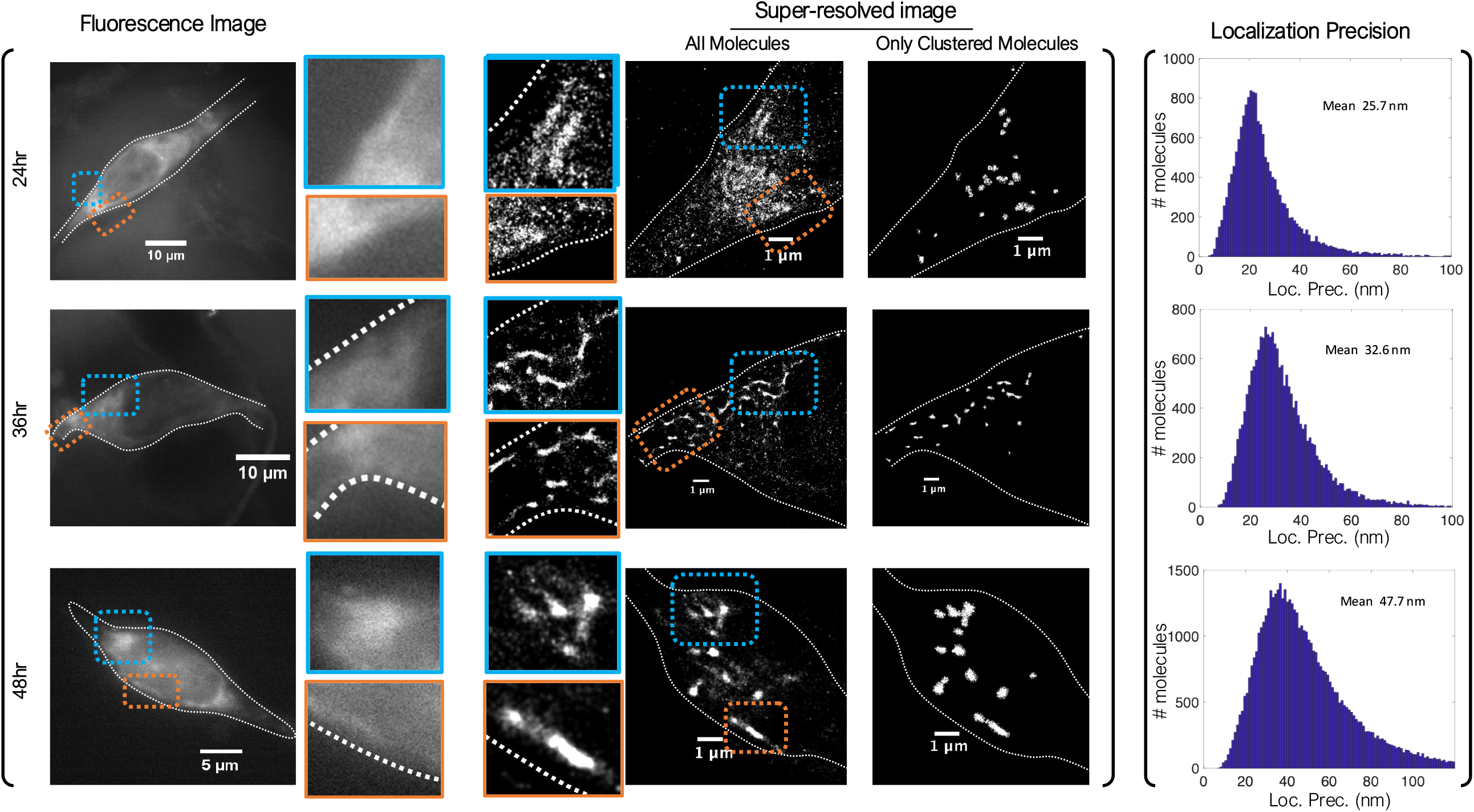
Super-resolution imaging at varying time-points (24 hrs, 36 hrs, 48 hrs): SMLM images (all molecules and only the clustered molecules) of fixed cells transfected with Dendra2-EboVGP showing distribution of single molecule and their clusters. Alongside enlarged images are also shown. Corresponding localization precision histogram showing a mean localization precision of in the range, 25 − 47*nm*.

### D. Biophysical parameter estimation

A better insight into the viral infection process is possible if the protein clusters are analyzed. Accordingly, single-molecule cluster analysis is carried out, and critical biophysical parameters are determined. The parameters include the size of cluster, the number of molecules per cluster, and the packing density of these viral proteins. Single molecule studies are carried that shows an average cluster area of 0.0337 *µm*^2^, 105 molecules per cluster and a density of 3351*mol*.*/µm*^2^ post 24 hrs of transfection (see, Fig. 5). This changes over time, with the cluster size (area) expanding to 0.1219 *µm*^2^ and a density of 5345 *mol*.*/µm*^2^ with an average of 642 molecules per cluster at 48 hrs. This is also evident from the average parameter plot, which shows a trend with all the parameters increasing proportionally with time. This indicates a large molecular fraction with time, suggesting increased participation of viral molecules in the clustering process.

**FIG. 5:**
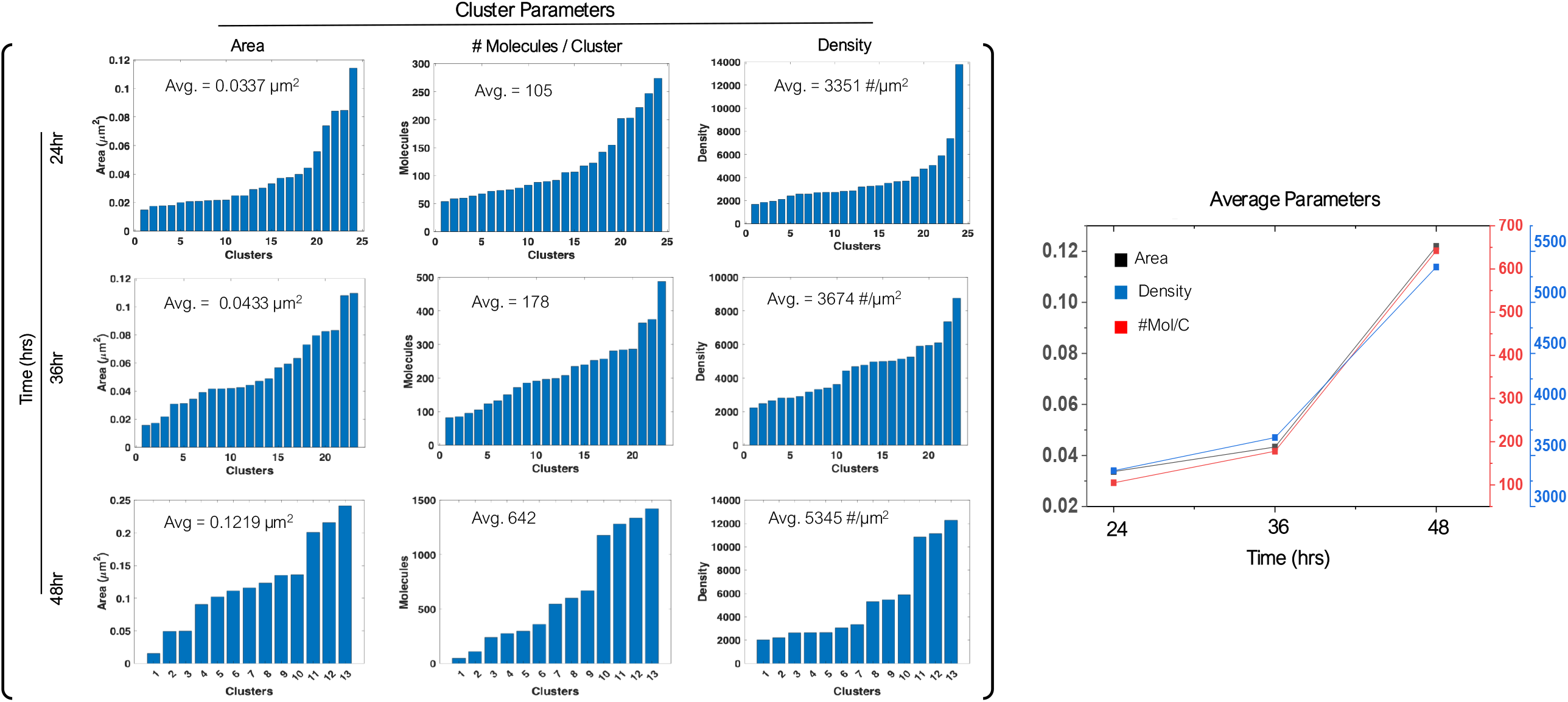
Cluster Parameter Estimation: Statistical analysis of single molecule data to estimate cluster parameters (cluster area, number of molecules per cluster and cluster density) for all time-points (24 hrs, 36 hrs and 48 hrs), post transfection. Plot of average cluster parameters showing the interdependence over time.

### E. Time-lapse super-resolution imaging

Time-lapse super-resolution imaging is carried out in a live transfected cell to understand the process of EboVGP clustering in a cell, which allows direct visualization of the process. Fig. 6 shows super-resolved images of a live cell observed over 1 hour, post 24 hrs of transfection. Alongside, the transmission image and fluorescence of the transfected cell are also shown. It is visually apparent from the time-lapse images (obtained at time-points, 0, 30, 60 mins) that the viral molecules undergo reorganization, especially during the early phases, i.e, just after 24 hrs of transfection. This ultimately gives rise to stable clusters as shown in Fig. 4.

**FIG. 6:**
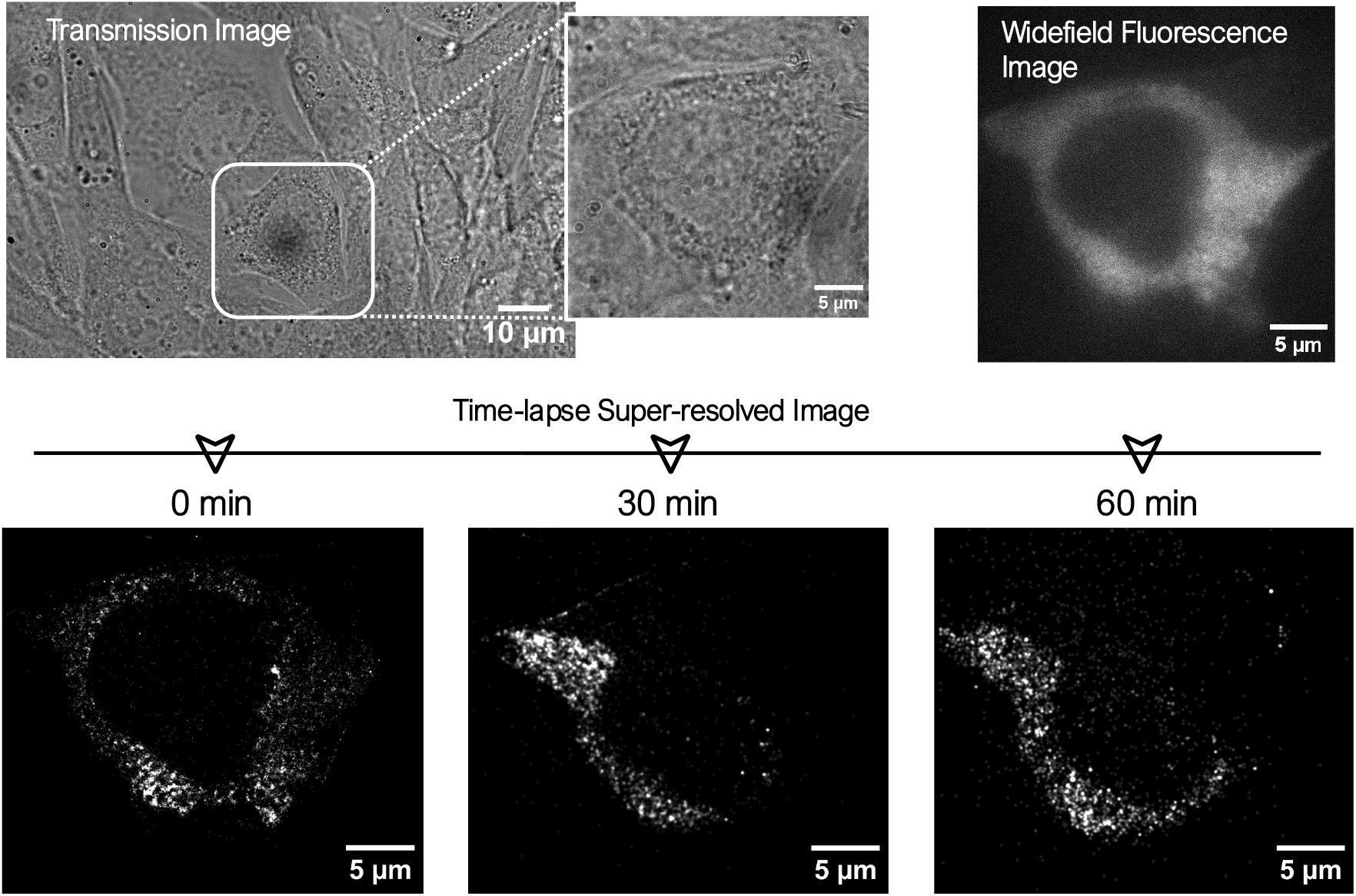
Time-lapse imaging of Dendra2-EboVGP, post 24 hrs of transfection: Single molecule data is recorded at an interval of 30 minutes for 1 hr, post 24 hrs of transfection. Alongside transmission image and diffraction-limited fluorescence microscopy image is also shown.

### F. Summary & Mechanism

Finally, the confocal and super-resolution microscopy study reveals the underlying mechanism that led to the formation of EboVGP clusters in a NIH3T3 cell, as pictorially represented in Fig. 7. Upon transfection of NIH3T3 cells, the lipo-complex containing pDendra2-EbGlyc enters the cell through endocytosis, followed by uncoating. Subsequently, it enters the nucleus where synthesis of mRNA takes place. *mRNA* is then carried to the ribosomes (located on the ER), followed by translation. Finally, this results in the expression of protein and its accumulation in the target organelle (plasma membrane) of the cell.

**FIG. 7:**
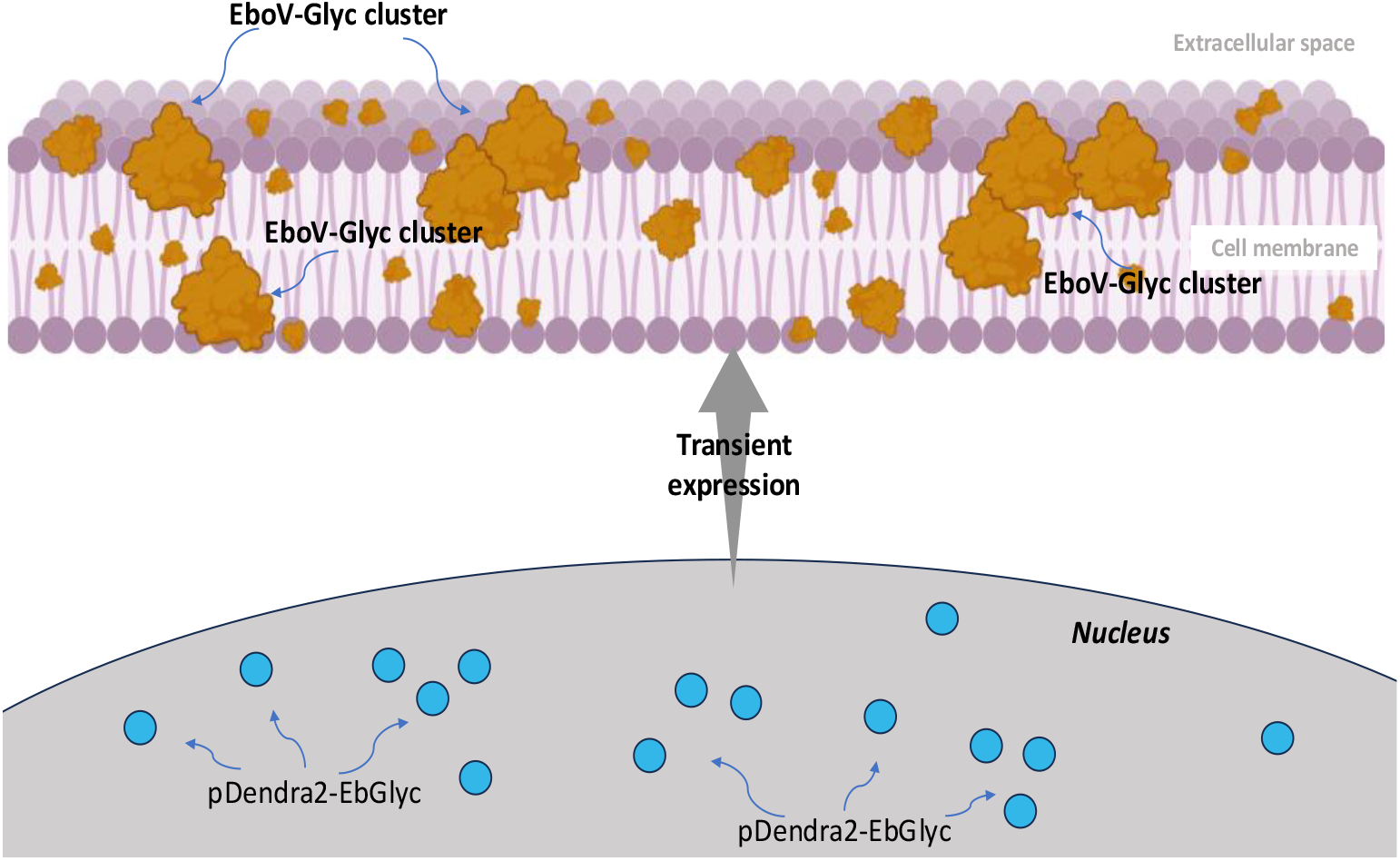
Dendra2-EbovGP Summary & Mechanism: Cartoon showing the mechanism of EboVGP, its clusters and localization to plasma-membrane of a cell, beginning from its processing (transcription, translation and others) and transient expression.

## Discussion & Conclusions

Understanding the viral infection mechanism and the underlying biological process in a cellular system at the single-molecule level is crucial for combating Ebola. The present study reports the dynamics of a single viral protein (EboVGP) and its clustering in a cellular system. This is made possible by single-molecule-based super-resolution microscopy that can resolve features down to a few tens of nanometers. The cell-based study reveals the details of the viral particles in the organelles of activity with the best possible resolution.

To facilitate SMLM-based super-resolution study and singlemolecule analysis, a recombinant photoactivable probe Dendra2-EbovGlyc (plasmid) is constructed by combining the protein-of-interest (EboVGP) and the marker protein (Dendra2) (see Fig. 1A,B). This is central to carrying out super-resolution imaging of transfected cells. The expressed protein showed appreciable transfection efficiency (4-5 transfected cells among 20 cells in the microscope field-of-view) and gave strong fluorescence when excited by blue light (fluorescence from non-activated molecules) (see, Fig. 1C). In addition, the proteins exhibit high brightness and signal-to-noise ratio that is crucial for detecting single molecule PSFs during superresolution imaging (*ex* : *em* = 561 *nm* : 579 *nm*). These qualities ensure detection of single viral protein molecules (Dendra2-EboVGP).

Our study and recent investigation suggest the dominant presence of viral protein in the plasma membrane of a cell (See Fig.2 and Fig. 3). This is evident from the localization studies post 24 hrs of transfection (Fig. 2), and from the prolonged study carried over 48 hrs post transfection (Fig. 3). The confocal studies and statistical analysis on transfected NIH3T3 cells indicate a high degree of colocalization of the viral protein (EboVGP) and the plasma-membrane. Other groups have also carried out similar studies. However, detailed studies of the expressed viral protein and its cluster are not possible using diffraction-limited microscopy (confocal). In addition, the analytical techniques can at best give indirect evidence of underlying biological processes and thus cannot be considered direct or conclusive.

To overcome this, we used super-resolution microscopy that can resolve fine features down to a single protein molecule and enable single molecule analysis. Here, we use single-molecule localization microscopy (SMLM) to show the location of activity of EboVGP in the cell plasma membrane, and formation of EboVGP clusters (see Fig. 4). These clusters have characteristic properties with specific sizes, number of molecules per cluster, and density (see Fig. 5). Surprisingly, the parameters show correlative behaviour with increase in cluster size (area), density, and the number of molecules per cluster with time (upto 48 hrs). This overall suggests an increase in the number of clustered molecules over time, i.e, more molecules participate in cluster formation. Finally, time-lapse imaging of live NIH3T3 cells transfected with recombinant probe Dendra2-EboVGlyc is carried out (see Fig. 6). The study shows a change in the number of molecules with time, indicating their migration. This is also an indication of single-molecule dynamics and its reorganization from one part of the plasma membrane to another during the initial stages of cluster formation. The whole process of cluster formation in the proposed transfection study is encapsulated in a cartoon (see, Fig. 7).

Overall, SMLM super-resolution microscopy has advantages over existing state-of-the-art confocal microscopy techniques. It has the potential to reveal details with resolution far below the diffraction limit of light and enables single-molecule analysis. This is seldom possible using existing methods. It is expected that the technique can reveal new information at the single-molecule level to understand the viral infection process in Ebola and related infectious diseases.

## III. METHODS

### A. Recombinant Plasmids

Recombinant plasmid Dendra2-EboVGlyc was generated by sub-cloning of Ebola Virus (Makona) Glycoprotein from pGL4.23 A82V 2014 EBOV + Mucin-Like Domain, a Gift from from Jeremy Luban and Pardis Sabeti [17] at SacI-BamH1 site of Dendra2-NS3 plasmid, from host lab [38]. The Ebola Virus (Makona) Glycoprotein sequence with Sac1-BamHI overhang was generated by PCR using Ebola glycoprotein specific oligonucleotide primers, EboVSacFP with Sac1 site as forward primer and EboVBamRP with Bam H1 site as reverse primer. The List and sequence of primers used are given in Table 1. PCR was performed using Phusion High-Fidelity DNA Polymerase (Thermo Fisher Scientific, India). The restriction enzymes used were purchased from Thermo Fisher Scientific, India. The Subcloning and transformation were performed using the standard protocol[**?**]. The recombinant plasmid Dendra2-EBoVGlyc was used to transfect NIH3T3 cells (mouse embryonic fibroblast cell line) to study the intracellular interaction of EboVGP.

**TABLE I:**
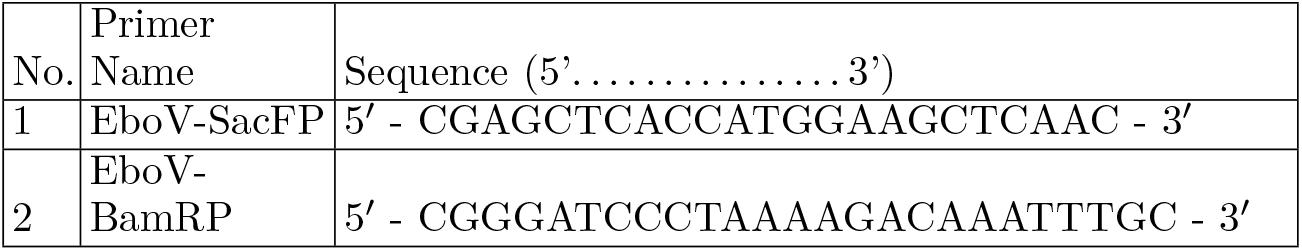
List of primers used for the construction of recombinant probe Dendra2-EboVGlyc.

### B. Cell Line and Transfection

The recombinant plasmid, Dendra2-EBoVGlyc was used to transfect NIH3T3 cells (mouse embryonic fibroblast cell line) (ATCC, USA) to study the interaction of the Ebola virus glycoprotein in cellular system. The cells, grown for 12h on a coverslip (No.0) (Blue star, India) in a 35mm dish with a density of 10^5^ cells/ml were transfected with 1 *µg* of recombinant plasmid DNA using Lipofectamine 3000 (Invitrogen, USA) according to manufactures protocol. The transfected dishes were incubated for 24 h. After 24 h of post-transfection, cells were washed twice with 1X PBS, and fixed with 3.7% (W/V) paraformaldehyde for 15 min. It was followed by washing with 1X PBS two times. Coverslip was removed from the 35mm dish and placed on a clean glass slide upside down with mounting media fluorosave (Thermofisher, USA). The fixed cells were made airproof by applying nail paint on the edges for long time preservation and further investigation. The same protocol was followed for 36 h and 48 h samples also.

### C. Image Acquisition Using Confocal Microscopy

The cells that express Dendra2-EboVGP are imaged on an Andor Dragonfly spinning disk confocal microscope(Bioimaging facility, Indian Institute of Science, Bangalore, India). Initially, transfected cells are identified using blue light on the eyepiece and the transfected cell is kept in the middle of the field of view. After that, the plasma membrane of the cell that is stained with Liss-Rhod-PE is imaged with the laser of wavelength 561 nm and the emission signal is collected from 572-616 nm subsequently, Dendra2-EboVGP was excited by a laser of wavelength 488nm, and the emitted fluorescence signal is collected within the spectral region of 502-540 nm. Along with fluorescence images, the brightfield image of the respective cell is also recorded. The fluorescence signal is collected using a 100x objective lens with a Z step size of 250nm. All the recorded images are deconvolved using Fusion software. The deconvolved images are further merged into a single 2D image by the maximum projection method, and their contrast is enhanced for better visualization.

### D. Single-Molecule Data Acquisition and Analysis

The single molecule data are acquired using lab-made Single Molecule Localization Microscopy, which consists of an Nikon Ti2E inverted microscope, fluorescence module, super-resolution module and detection module. The fluorescence module, which consists of blue light of wavelength 470-490nm, helps to identify properly transfected cells in the sample. Once the transfected cell is identified, the microscope is switched to super-resolution mode that consist activation laser (405 nm, OBIS) and readout laser(561nm, Oxxius). These two lasers are combined using a dichroic mirror(DMLP425R, Thorlabs) and directed towards the back port of the microscope and focused at the backaperture of 100x objective lens using Electrically Tunable Lens(ETL). The ETL enable the user to change the field of view by underfilling and overfilling the backaperture of the objective lens. The single molecules are stochastically activated and excited with the laser and emits burst of photons. The emitted fluorescence photons are recorded using a detection module. The detection module contains an EMCCD(Andor, iXon ultra 897) camera, filter tube and a 4f system which is placed between camera and the microscope. The 4f system consists of two biconvex lenses of focal length 200mm and 75mm, and it gives a magnification of 2.66. This provides the overall system magnification to be 266X. The physical pixel size of the camera is 16*µm* and after 266X times magnification the effective pixel size(q) becomes 60nm (16*µm/*266 = 0.06*µm*) that ensures the oversampling of single molecule PSF. The filter tube contains two notch filters(NF03-405E-25 and NF03-561E-25) to eliminate activation and excitation laser and a long pass filter(LP02-561RE-25) to allow the fluorescence signal. The EMCCD camera sensor is cooled to − 88^*°*^*C*, to reduce thermal noise. The exposure time and EM gain are set up at 30ms and 270 respectively. The data is recorded in ‘.tif’ file format and processed using MATLAB programs. The background noise is removed using rolling ball algorithm and single-molecule spots are identified in each frame using a threshold value on number of photons. Further,the detected single-molecule spots are fitted with a 2D Gaussian function. The centroid, number of photons (N), background noise (b), and standard deviation (s) of the 2D Gaussian function of each fitted molecular spot are estimated from the fit parameters. In addition to that, localization precision (*σ*_*lp*_) of each molecule is calculated using Thompsons’ relation [42], 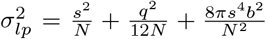 Finally, a super-resolution image is reconstructed by representing all the detected single molecules as a 2D Gaussian function whose standard deviation is taken as molecules’ localization precision.

### E. Colocalization Analysis

The colocalization analysis is performed using JACoP ImageJ plugin [15]. Initially, the transfected cell is masked among other cells in the Dendra2-EboVGP. The same mask is used for the plasma membrane channel. Later, using the masked images, Pearson Coefficient, and Overlap Coefficient between Dendra2-EboVGP and Plasma membrane are calculated.

### F. Single-Molecule Cluster Analysis

Dendra2-EboVGP molecule clusters are identified using DBSCAN (Density Based Spatial Clustering of Applications with Noise) algorithm. The Matlab built-in function *dbscan* is used. The function requires three input parameters such as X and Y coordinates of all the molecules as an array, minimum number of molecules in a cluster, and maximum distance between two molecules to be considered for clustering. The minimum number of molecules in a cluster is considered as 50 and the maximum Euclidean distance between the molecules is considered as 2 pixels (120nm). Once the molecules are grouped into clusters, the area of each cluster, the number of molecules per cluster, and the density(number of molecules per unit area) are calculated. The entire cluster analysis is performed in MATLAB R2021a.

### G. Time-Lapse Imaging

The cell sample is prepared in a 35mm dish with a bottom coverslip. 24 hours post-transfection, the cell culture media in the dish is removed and 1 ml of live cell imaging solution(Gibco, A59688DJ) is added. The imaging buffer minimize the photo bleaching, reduces the background fluorescence and maintain physiological condition of cells. Single-Molecule Localization Microscopy is used to perform Time-Lapse imaging. The lasers are operated at low power to reduce to photo toxicity (activation laser power is 8 *µW* and excitation laser power is kept at 18 *mW*). A total of 5000 frames with the exposure time of 40 *ms* are recorded for each time point, and the experiment is performed for a period of 1 hour with an interval of 30 minutes. The data are recorded in the frame transfer mode to minimize the data acquisition time and this also helps the sample to be exposed by laser for lesser time.

## Acknowledgements

The authors thank Dr. Subhra Mandal (University of Nebraska - Lincoln, USA) for going through the manuscript and providing valuable inputs.

## Author Contributions

PPM conceived the idea. JMV, AS, NP, PPM carried out the experiments. JMV, AS, NP, PPM prepared the samples. AS, NP and SS carried out the analysis. PPM wrote the paper by taking inputs from all the authors.

## Notes

### Competing Interest Statement

The authors have declared no competing interest.

